# Genomic transcription factor binding site selection is edited by the chromatin remodeling factor CHD4

**DOI:** 10.1101/2023.03.02.530867

**Authors:** Mika Saotome, Deepak Balakrishnan Poduval, Sara A. Grimm, Aerica Nagornyuk, Sakuntha Gunarathna, Takashi Shimbo, Paul A. Wade, Motoki Takaku

## Abstract

Biologically precise enhancer licensing by lineage-determining transcription factors enables activation of transcripts appropriate to biological demand and prevents deleterious gene activation. This essential process is challenged by the millions of matches to most transcription factor binding motifs present in many eukaryotic genomes, leading to questions about how transcription factors achieve the exquisite specificity required. The importance of chromatin remodeling factors to enhancer activation is highlighted by their frequent mutation in developmental disorders and in cancer. Here we determine the roles of CHD4 to enhancer licensing and maintenance in breast cancer cells and during cellular reprogramming. In unchallenged basal breast cancer cells, CHD4 modulates chromatin accessibility at transcription factor binding sites; its depletion leads to altered motif scanning and redistribution of transcription factors to sites not previously occupied. During GATA3-mediated cellular reprogramming, CHD4 activity is necessary to prevent inappropriate chromatin opening and enhancer licensing. Mechanistically, CHD4 competes with transcription factor-DNA interaction by promoting nucleosome positioning over binding motifs. We propose that CHD4 acts as a chromatin proof-reading enzyme that prevents inappropriate gene expression by editing binding site selection by transcription factors.

## Introduction

During the cell fate transitions integral to development or transcription factor-dependent cellular reprogramming, lineage-determining transcription factors (TFs) must contend with chromatin to nucleate active enhancers. Some transcription factors such as pioneer factors utilize their intrinsic ability to bind nucleosomal DNA in closed chromatin as a first step in the induction of chromatin opening ^1, 2^. Establishment of an active enhancer is hypothesized to be a multi-step process involving alterations to local chromatin by chromatin remodeling enzymes, histone replacement, and editing of histone and DNA modification ^3, 4^. All downstream processes flow from the initial event – recognition of DNA sequence by a sequence-specific DNA binding transcription factor. In most cases, TFs are present on the order of thousands to tens of thousands of molecules per cell. In contrast, many transcription factor motifs are present on the order of millions of copies per genome. How TFs are directed to activate enhancers in all the correct locations and only the correct locations is critical, as both failure to activate appropriate sets of genes as well as inappropriate gene activation is typically deleterious to the biological program. Cooperative binding by multiple TFs, partial motif recognition, nucleosome positioning, and chromatin remodeling factors are thought to be involved in selective enhancer formation ^3, 5–7^. However, the molecular mechanisms remain elusive.

CHD4 (Chromodomain Helicase DNA Binding Protein 4) is a catalytic core component of the NuRD (Nucleosome Remodeling and Deacetylase) chromatin remodeling complex and is known to regulate gene expression and DNA damage responses ^8, 9^.

CHD4 is involved in multiple developmental processes including neural development and cardiac development ^10–12^. Similar to other chromatin remodeling factors, genetic and epigenetic data in cancer patients detected frequent alterations in the CHD4 gene suggesting key roles of CHD4 during tumorigenesis and tumor progression ^8, 13–16^.

Because the NuRD complex contains histone deacetylases such as HDAC1 or HDAC2, it has been thought to act as a gene silencer ^17–20^. However, the genomic distribution of CHD4 is predominantly enriched at promoters or open chromatin regions, and the CHD4 function appears to be cell context specific ^21–24^.

In this study, we measured the impacts of CHD4 depletion in MDA-MB-231 basal breast cancer cells. Characterization of chromatin features by multi-omics methodologies revealed differential impacts of CHD4 knockdown in steady-state cells vs cellular reprogramming processes. CHD4 knockdown in steady-state MDA-MB-231 cells tends to increase chromatin accessibility at promoter-distal regions. Mechanistically, AP1 family transcription factors are involved in altered chromatin binding accessibility and transcription. During the GATA3-induced MET cell reprogramming processes ^25^, CHD4 maintains chromatin architecture mainly at intergenic regions. In the absence of CHD4, chromatin sensitivity was increased at closed chromatin, especially at GATA3 binding peaks. Abnormal chromatin opening led to increased expression of genes unrelated to MET. High-resolution nucleosome mapping suggested that CHD4 prevents unnecessary gene expression by mediating nucleosome formation over the GATA3 binding sites. These results demonstrate that CHD4, and by extension NuRD complex, monitors transcription factor/chromatin interactions and modulates gene regulation by proofreading transcription factor site selection for enhancer nucleation.

## Results

### CHD4 depletion mediates altered chromatin accessibility and gene expression

To explore the range of chromatin features regulated by CHD4 in somatic cells, we depleted it in MDA-MB-231 basal breast cancer cells using shRNA (Supplementary Figure 1a) followed by genome-wide analysis of chromatin accessibility using ATAC-seq ^26, 27^. The physical location of most ATAC-seq peaks in the genome was unchanged, while approximately 1/3 of detected peaks were either lost or gained following CHD4 depletion (Figure 1a - 1b). Comparisons at the level of individual biological replicates indicated that loss of CHD4 rather than inherent variability drives the outcome of this comparison (Supplementary Figure 1b - 1c).

**Figure 1.**
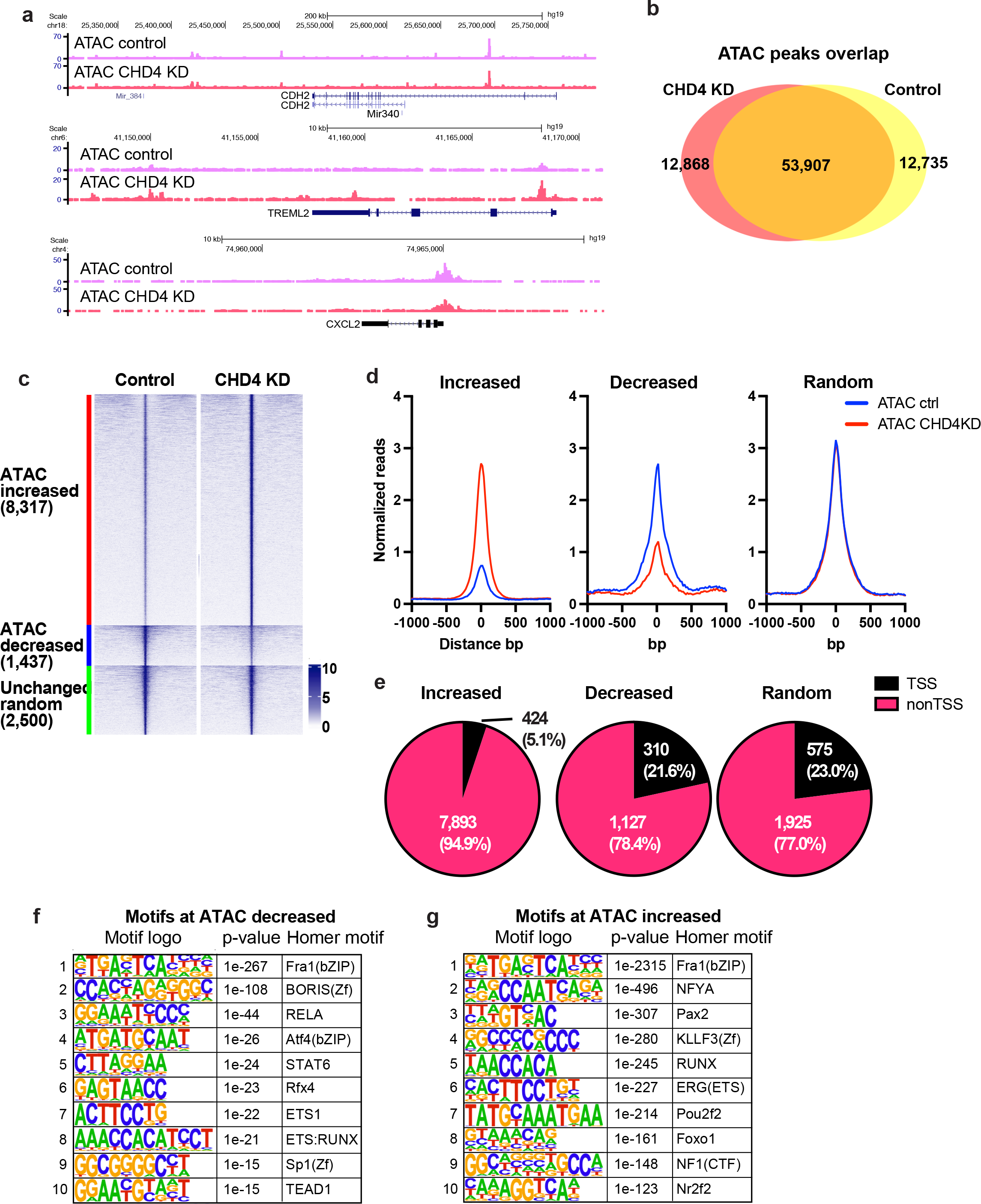
CHD4 modulates chromatin accessibility at promoters in MDA-MB-231 **(a)** Genome browser tracks showing ATAC-seq signals in control or CHD4 knockdown (KD) cells. Unchanged, increased, and decreased regions are selected. **(b)** Venn diagram showing the ATAC-seq peak overlap between control or CHD4 KD cells. **(c)** edgeR differential ATAC-seq peak analysis in CHD4 KD cells. FDR < 0.01 and |fold change| > 2 are applied to define differential peaks. Heatmap showing ATAC-seq signals at increased, decreased, or randomly selected unchanged ATAC-seq peaks. **(d)** Metaplot showing normalized ATAC-seq reads/peak at differential peaks. **(e)** Pie chart showing the frequency of TSS and non-TSS peaks in each peak group. **(f-g)** HOMER de novo motif analysis. Decreased (f) or increased ATAC-seq peaks (g) are used as input.

In addition to loss or gain of individual peaks, we analyzed the data to assess whether individual loci had changes in the level of accessibility with or without a change in location. Somewhat paradoxically, most altered ATAC peaks exhibit an increase in transposition following depletion of CHD4 (Figure 1c - 1d). The loci with increased accessibility were largely confined to peaks located greater than 1 kb from an annotated transcription start site (TSS) and this distribution differed from random (p-value = 0.00001, Chi Square Test) (Figure 1e, Supplementary Figure 1d). Peaks with decreased accessibility were associated with transcription start sites at a frequency similar to a random peak set (Figure 1e). These results suggest that CHD4 may have a different impact on chromatin architecture at promoters than at distal regulatory elements.

We asked whether the alteration in peak intensity reflected changes in the binding behavior of individual transcription factors by assessing the enrichment of these loci for known transcription factor binding motifs using HOMER ^28^. Surprisingly, there was considerable overlap in the binding motifs enriched in the peaks with increased accessibility and the motifs enriched in the peaks with decreased accessibility. Motifs for the AP1 family, the ETS family and the RUNX family were present in both enriched motif sets, with an AP1 motif being the most enriched in both sets (Figure 1f - 1g). To further access AP1 motif enrichment, we performed the HOMER motif enrichment analysis using the gained (12,868 ATAC-seq peaks, uniquely observed in CHD4 KD cells) and lost peaks (12,735 ATAC-seq peaks, only observed in control cells) defined by peak overlap analysis shown in Figure 1b. Similarly, motifs for the AP1 family members were significantly enriched at both gained and lost peaks (Supplementary Figure 1e - 1f). This outcome was unexpected, as it suggested that loss of CHD4 leads to an alteration in the competition between transcription factors and nucleosomes for specific DNA.

To assess whether individual transcription factors were, in fact, relocalized in the genome, we performed ChIP-seq for AP1 family members, JUNB, FRA1, and ATF3 with and without depletion of CHD4. Consistent with the ATAC-seq data, all tested AP1 family members showed thousands of differential bindings upon CHD4 depletion (Figure 2a). For the case of JUNB, 2,190 increased and 3,301 decreased peaks were observed (Figure 2b - 2c). Slightly smaller numbers of differential binding peaks were observed for FRA1 and ATF3 (Supplementary Figure 2a - 2b). Metaplot analysis at ATAC-seq differential peaks suggested that increased JUNB and FRA1 binding was associated with increased chromatin accessibility, while decreased chromatin accessibility was associated with reduced ATF3 binding. To ask whether these alterations in chromatin were associated with CHD4 binding, we performed ChIP-seq for CHD4 in MDA-MB-231 cells. CDH4 peaks were frequently observed at open chromatin regions (Figure 2d), and approximately 80% CHD4 peaks (13,419 peaks out of 16,780 peaks) were overlapped with ATAC-seq peaks in MDA-MB-231 cells (Figure 2e).

**Figure 2.**
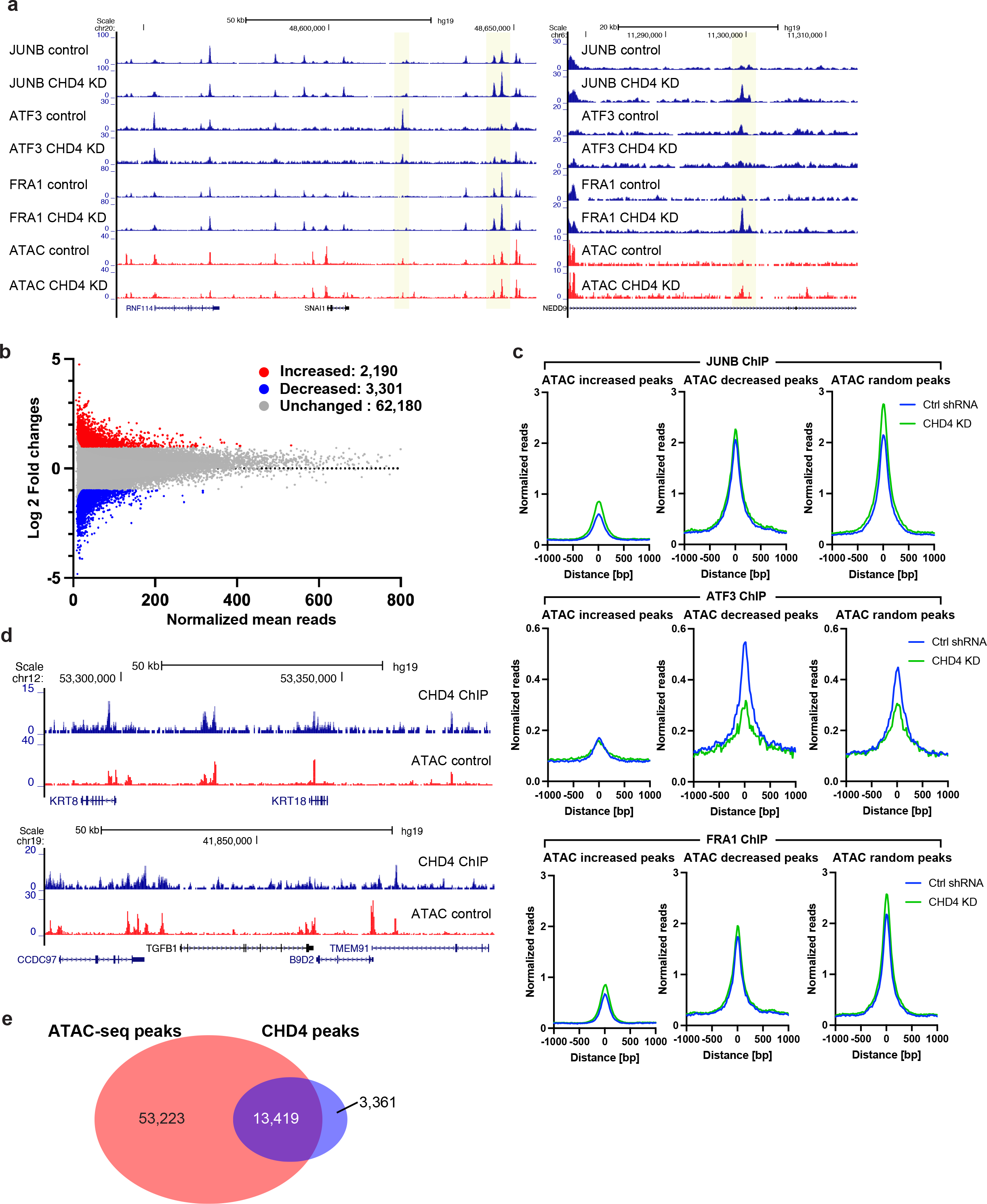
AP1 family transcription factors are redistributed following CHD4 depletion **(a)** Genome browser tracks showing JUNB, ATF3, and FRA1 ChIP-seq data. Differential peaks between control and CHD4 KD data are highlighted in yellow. ATAC-seq data are also shown as a reference for open chromatin regions. **(b)** Scatter plot showing increased (red), decreased (blue), and unchanged (grey) peaks. FDR < 0.01 and |fold change| > 2 are applied to define differential peaks by edgeR. **(c)** Metaplot showing normalized ChIP-seq reads/peak at ATAC-seq differential peaks. JUNB (top). ATF3 (middle), and FRA1 (bottom) ChIP-seq signals in control (blue) or CHD4 KD cells (green) are plotted. **(d)** Genome browser tracks showing the frequent overlap between CHD4 and ATAC-seq peaks. CHD4 ChIP-seq were performed in control MDA-MB-231 cells. **(e)** Venn diagram showing the overlap between ATAC-seq peaks and CHD4 peaks.

Differential chromatin accessibility and redistribution of multiple AP1 family members upon CHD4 depletion suggested that CHD4 is important for target site selection by transcription factors. AP-1 proteins have a high affinity to the palindromic sequence 5’- TGA G/C TCA-3’, but transcription factors are known to possess nonspecific or non-consensus DNA binding. Therefore, CHD4 might regulate motif scanning activities of transcription factors. In the HOMER database, we found more than 1 million AP-1 potential binding sites. We first tested if we could detect consensus motif scanning from our JUNB and FRA1 ChIP-seq data. For this analysis, we excluded the observed ChIP-seq peaks to minimize the bias in the ChIP-seq data resulting from motif-independent binding or indirect binding caused by cell fixation (Figure 3a). Both JUNB and FRA1 showed significantly stronger enrichment at the consensus motif containing loci compared to randomly selected genomic regions (Figure 3b - 3c), consistent with motif scanning. When CHD4 was depleted, JUNB and FRA1 binding signals were decreased at the consensus motif sites (Figure 3d - 3e), suggesting that CHD4 modulates motif scanning activities of JUNB and FRA1, potentially permitting a more promiscuous scanning process.

**Figure 3.**
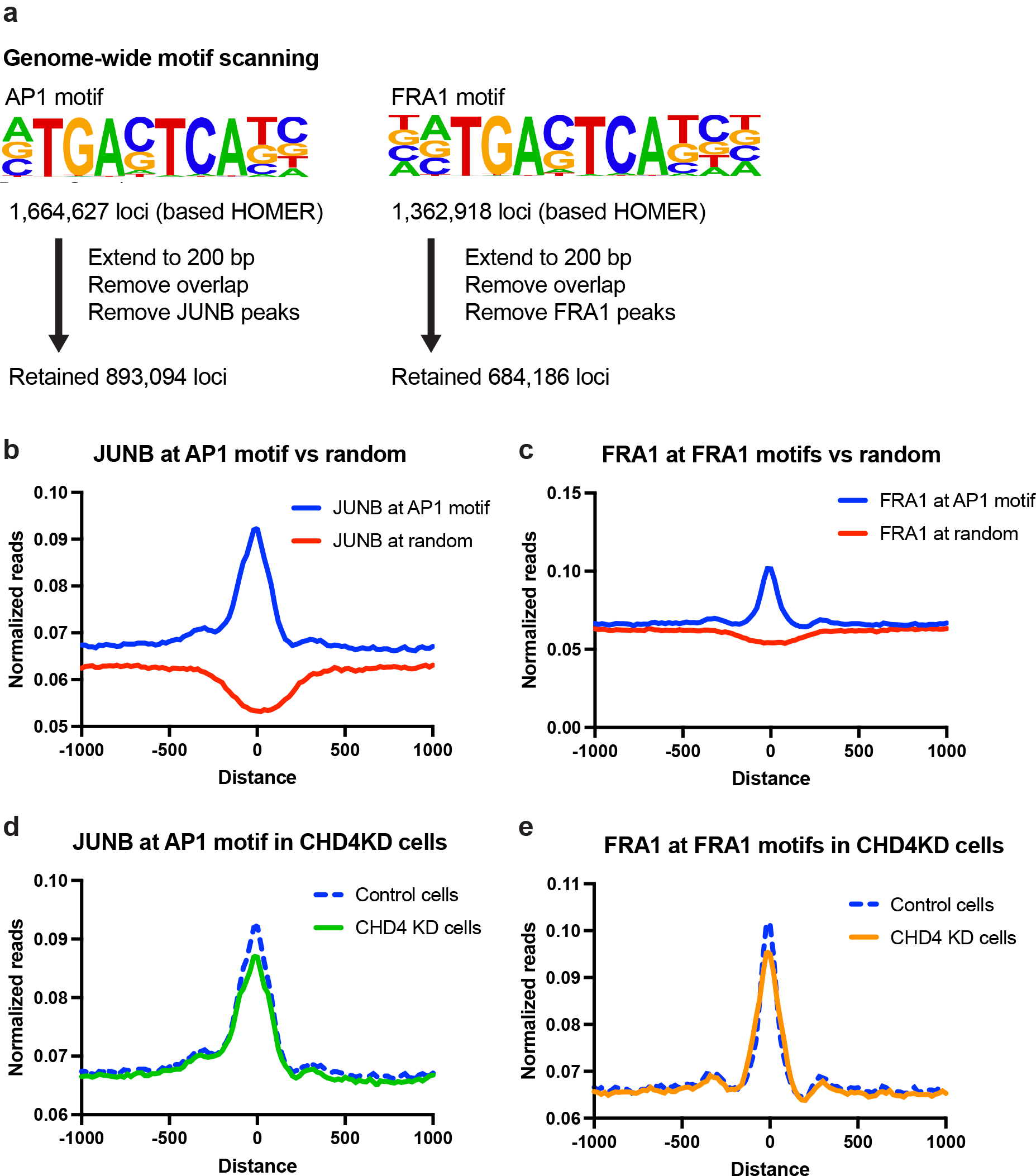
Genome-wide motif scanning of AP1 family proteins is affected by CHD4 knockdown **(a)** Scheme for defining potential binding target loci. HOMER AP1 or FRA1 motif-containing sites are obtained from the HOMER database. For the downstream analysis, each consensus motif locus was extended to 200 bp, and the overlaps within the consensus motif loci and with the observed JUNB or FRA1 ChIP-seq peaks were removed. The retained loci were used for detecting motif scanning activities of JUNB and FRA1. **(b)** Metaplot showing JUNB ChIP-seq signals at AP1 consensus motif (blue) or randomly selected genomic (red) regions. **(c)** Metaplot showing FRA1 ChIP-seq signals at FRA1 consensus motif (blue) or randomly selected genomic (red) regions. **(d-e)** Metaplot showing JUNB (d) or FRA1 (e) ChIP-seq signals at consensus motif regions in control (blue) or CHD4 KD (green) cells.

To examine the functional consequences to gene expression of CHD4 depletion, we performed RNA-seq. FDR < 0.05 and fold change criteria (|fold change| > 1.5) were used to identify significantly altered transcripts. We identified a large percentage of the transcriptome with significant alterations in steady state transcript level – 1880 genes were upregulated and 1072 genes were downregulated (Figure 4a). The number of genes altered in steady state transcript level was also skewed towards gene activation, but not to the extent as alterations in transposition. Integration of ATAC data and gene expression indicated that genes associated with increased transposition had higher steady state transcript levels following CHD4 depletion while genes with decreased accessibility had lower levels of transcript (Figure 4b). Gene ontology analysis suggested that altered transcript levels were enriched at genes involved in plasma membrane processes and interaction with the extracellular space (Figure 4c).

**Figure 4.**
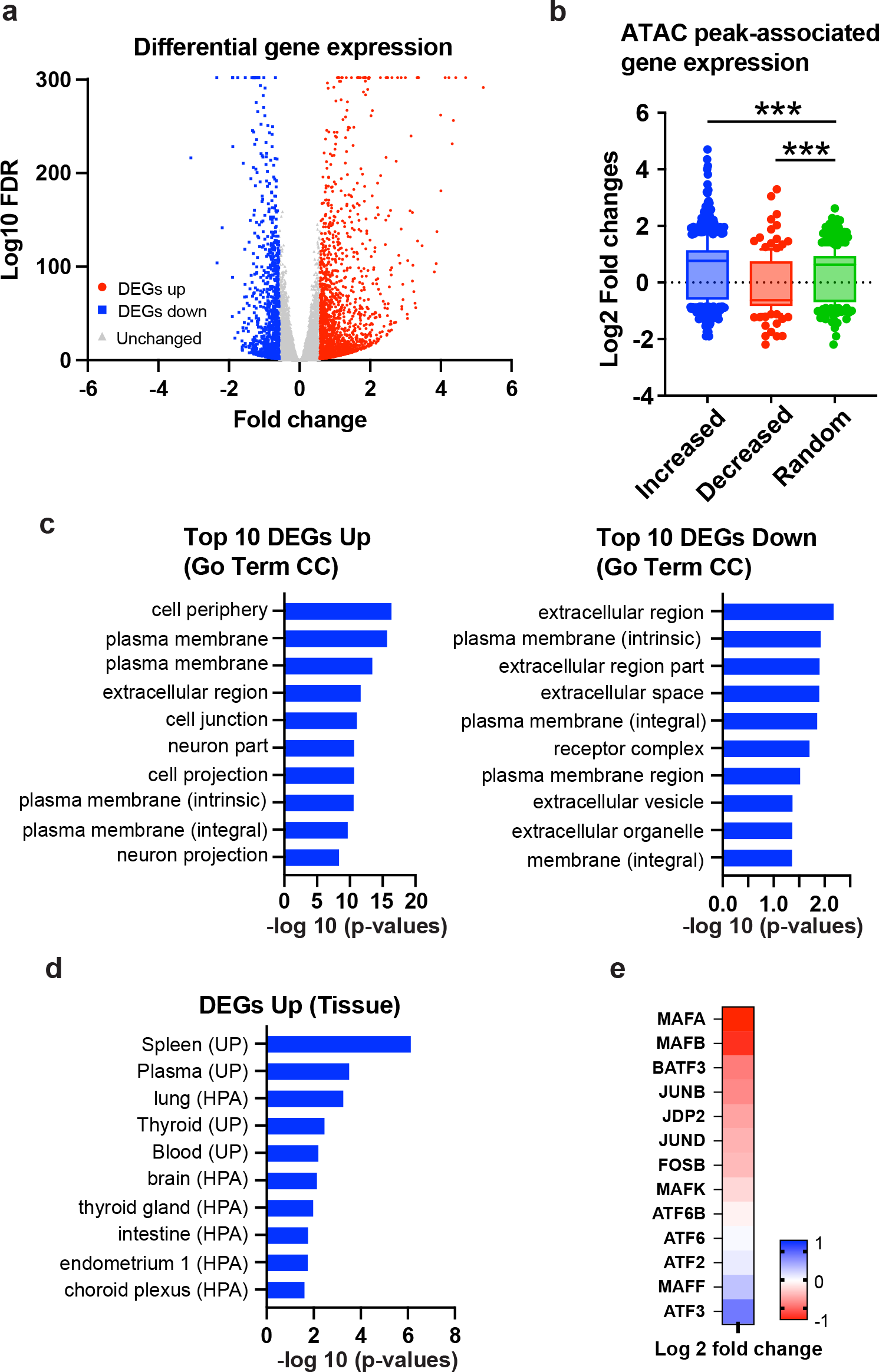
CHD4 knockdown results in aberrant gene expression unrelated to breast cancer program **(a)** Volcano plot showing differential gene expression upon CHD4 depletion. FDR < 0.05 and |fold change| > 1.5 were applied to define differentially expressed genes. **(b)** Gene expression of ATAC-seq differential peak associated genes. Box plots show log2 fold changes from each peak group. Increased, decreased, and randomly selected unchanged peaks are assigned to closest genes. **(c)** Pathway enrichment analysis. Most significantly altered genes (fold change +/- 2) were used for Go term Cellular Component (CC) analy- sis by DAVID. Top 10 pathways are indicated. **(d)** Functional annotation of up-regulated genes in CHD4 KD cells. DAVID tissue expression annotation was performed using up-regulated genes. **(e)** Gene expression of AP1 family proteins. ATAC-seq differential peak associated genes. Heatmap shows log2 fold changes in CHD4 KD cells compared to control shRNA condition.

Somewhat surprisingly, upregulated transcripts were associated with tissues other than breast (Figure 4d), suggesting that loss of CHD4 leads to loss of cell-type specificity in the transcriptional program. Direct inspection of the RNA-seq data revealed a large number of transcription factors that change in steady state transcript level, including AP1 family members (Figure 4e).

### CHD4 antagonizes enhancer formation by GATA3

We observed alterations in the dynamic competition between nucleosomes and transcription factors for DNA upon depletion of CHD4. However, we also observed changes in expression of transcription factor family members, potentially complicating analysis. Therefore, we moved to a more defined system with a temporal component, transcription-factor dependent cellular reprogramming. We established a doxycycline-inducible GATA3 expression system in MDA-MB-231 mesenchymal breast cancer cells. We selected a cell clone that shows minimal GATA3 expression in the absence of doxycycline (hereafter DOX) but expresses biologically relevant amounts of GATA3 protein upon DOX treatment. In this cell clone, GATA3 protein expression was observed within 3 hours after addition of DOX to media and was saturated by 12 hours (Supplementary Figure 3a). Stable GATA3 expression was observed for at least 48 hours after DOX induction.

We collected ATAC-seq, CHD4 and GATA3 ChIP-seq and RNA-seq 12 hours after DOX treatment to characterize GATA3 and CHD4-dependent changes in chromatin architecture and gene expression. By comparing transposase accessibility at GATA3 peaks before and after GATA3 induction, we characterized three predominant types of loci: loci that transition from inaccessible to accessible (newly accessible), loci where GATA3 binding occurs within transposase accessible chromatin (constitutively accessible) and loci where GATA3 binds to inaccessible chromatin that remains inaccessible (constitutively inaccessible) following GATA3 expression (Figure 5a - 5b, green curves). Overlap between CHD4 and GATA3 at newly accessible and constitutively accessible GATA3 peaks was substantial (Supplementary Figure 3b - 3c). RNA-seq analysis revealed that 926 genes were altered in steady state abundance following GATA3 expression, with slightly more genes being activated (524) than decreased (402) (Figure 5c). Consistent with previous reports ^25^, transcripts with altered levels were enriched in categories involved in mesenchymal to epithelial transition (Supplementary Figure 3d).

**Figure 5.**
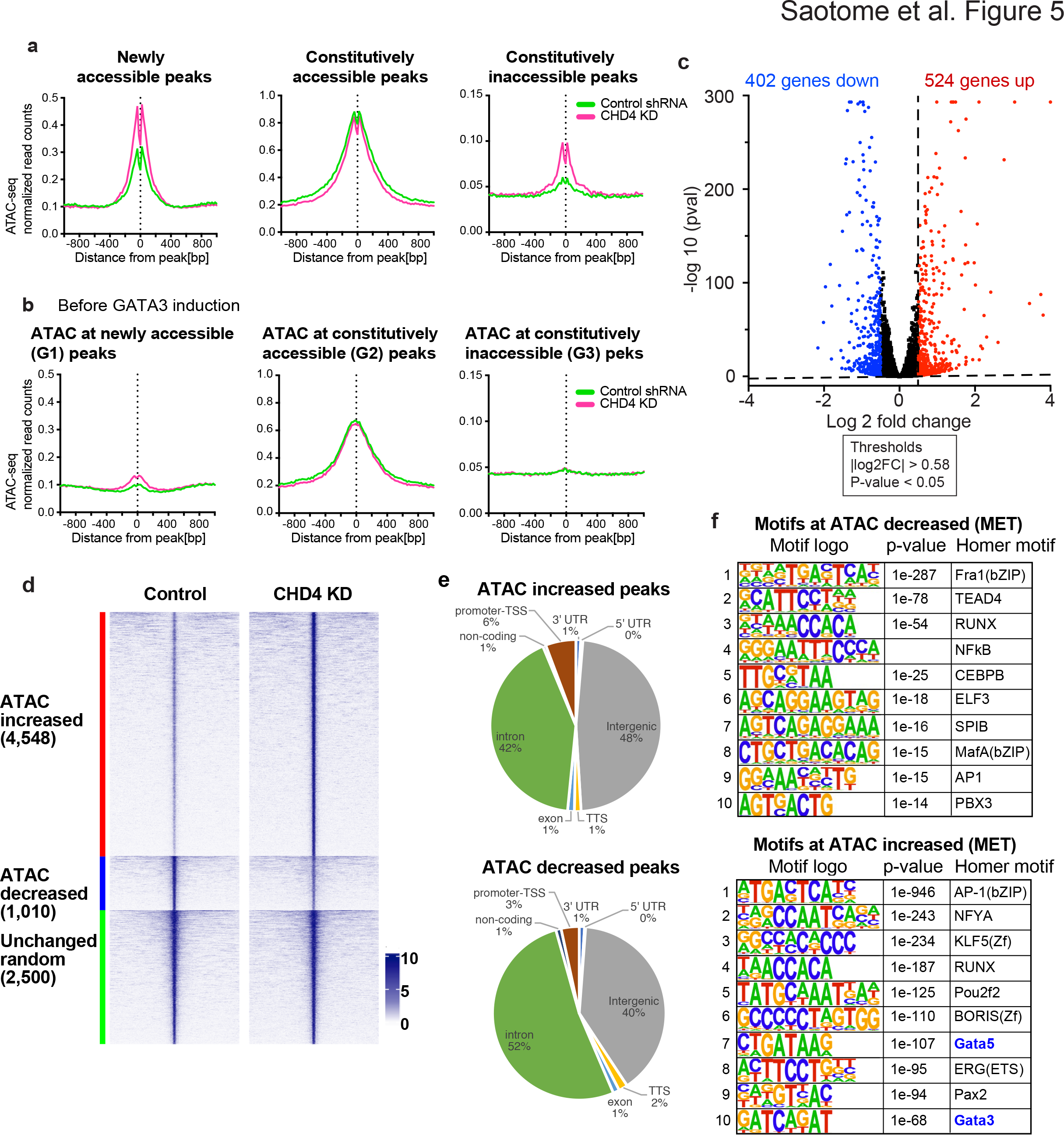
CHD4 knockdown leads to abnormal chromatin opening **(a-b)** Metaplots showing ATAC-seq signals in the control shRNA or CHD4 shRNA transduced cells. Averaged ATAC-seq signals before (b) or after (a) GATA3 expression are plotted in each peak group. **(c)** Volcano plot showing differential gene expression 12 hours after GATA3 expression. Up- and down-regulated genes (FDR < 0.05, |log2 (fold change)| >0.5) are highlighted in red and blue, respectively. **(d)** ATAC-seq differential peak analysis upon CHD4 knockdown in MET. RNAs were collected 12 hours after GATA3 expression. FDR < 0.01 and |fold change| > 2 are applied to define differential peaks. Heatmap shows ATAC-seq signal intensity in control or CHD4 KD cells at increased, decreased, and randomly selected unchanged peaks. ATAC-seq signals at increased, decreased, or randomly selected unchanged ATAC-seq peaks. (e) Pie chart showing peak annotation defined by HOMER. Increased or decreased ATAC-seq peaks are classified into 8 peak categories. (f) HOMER de novo motif analysis. Decreased (top) or increased (bottom) ATAC-seq peaks upon CHD4 KD in MET condition are used as input.

When CHD4 was depleted in this system, we observed striking alterations in several features. Of loci exhibiting significant alterations in ATAC sensitivity, 82% (4548 of 5558) demonstrated an increase in accessibility (Figure 5d), similar to control MB-MDA- 231 cells (Figure 1c). Unlike the case in control cells where increased accessibility was overwhelmingly at distal elements and decreased accessibility was somewhat more balanced, changes in transposition in the reprogramming system were overwhelmingly found distant from annotated transcription start sites (Figure 5e). Globally, altered ATAC loci were enriched in AP1 and RUNX motifs, with GATA motifs enriched at loci that gain accessibility (Figure 5f). When we focused analysis on loci with GATA3 peaks, we found that depletion of CHD4 led to a substantial increase in accessibility at newly accessible peaks where GATA3 binding leads to enhancer licensing (Figure 5a). Surprisingly, peaks where GATA3 fails to induce chromatin opening in the presence of CHD4 frequently display increased accessibility in its absence (Figure 5a), suggesting that CHD4 acts to oppose the chromatin-opening ability of GATA3.

RNA-seq performed following CHD4 knockdown revealed inappropriate gene expression at loci linked to constitutively inaccessible sites, exemplar genes are depicted in Figure 6a and Supplementary Figure 4. At such loci, GATA3 binding is not associated with transposase accessibility in the presence of CHD4, in its absence accessibility is evident along with increased transcript level. Gene ontology analysis revealed that the up-regulated genes associated with the constitutively inaccessible peaks were significantly enriched with tissue-specific genes related to GATA3 function in cell contexts other than breast (Figure 6b). For instance, GATA3 is known to be important for trophoblast differentiation and placental development ^29, 30^. Placenta related genes such as VSTM5, NDNF, HAPLN1 were up-regulated in the CHD4 knockdown cells (Figure 5a, Supplementary Figure 4).

**Figure 6.**
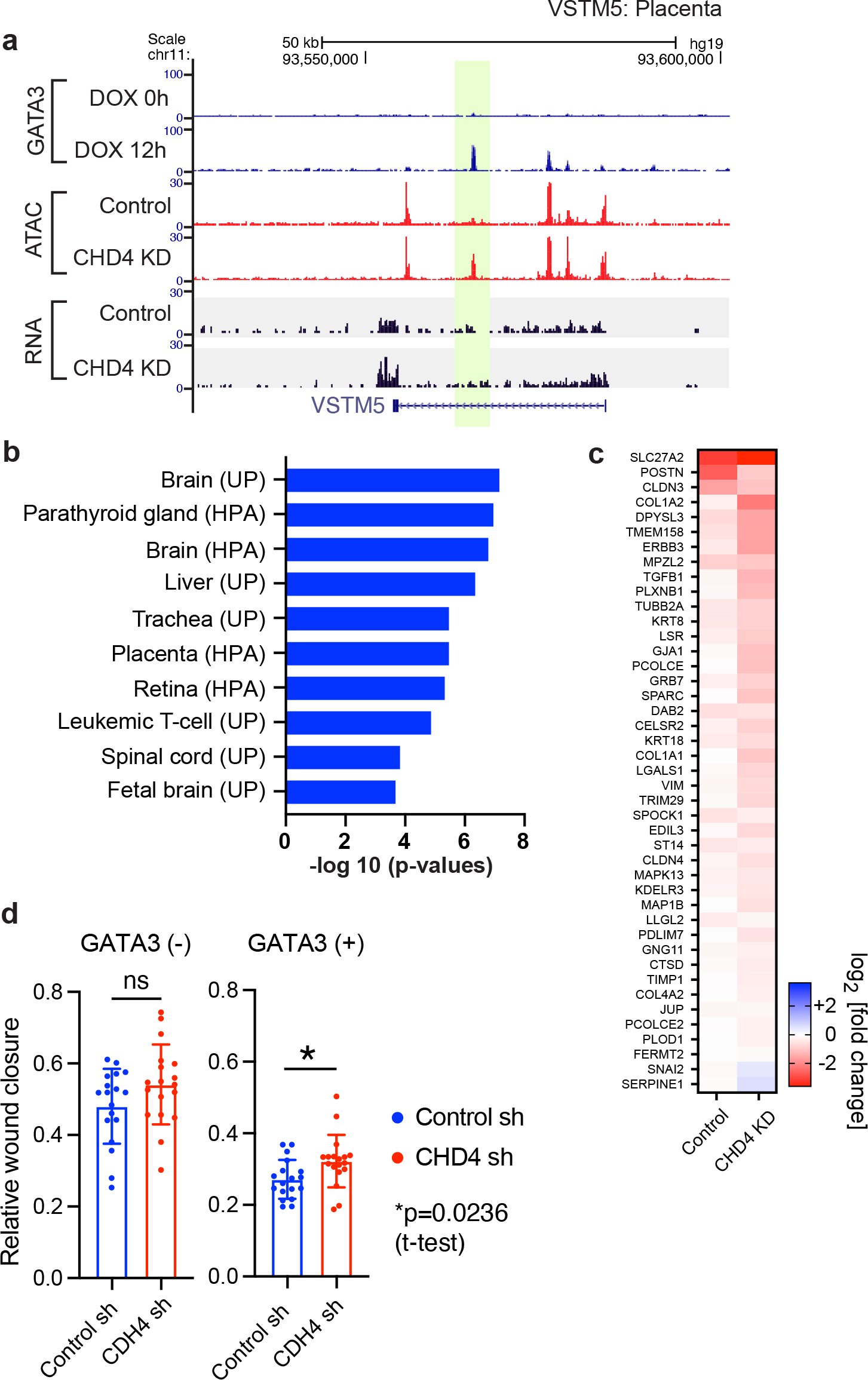
CHD4 regulates enhancer activities during MET cell reprogramming **(a)** An example of aberrant gene expression at a constitutively inaccessible GATA3 peak. Genome browser tracks show an example of up-regulated genes associated with the constitutively inaccessible GATA3 peaks. ATAC-seq and RNA-seq was performed 12 hours after GATA3 induction. The de novo open chromatin site upon CHD4 KD is highlighted in yellow. VSTM5 is known to be expressed in brain and placenta. **(b)** Functional annotation of the constitutively inaccessible peak associated genes. DAVID tissue expression annotation was performed using the up-regulated genes that are associated (+/- 100 kbp) with the constitutively inaccessible GATA3 peaks. **(c)** Heatmap showing expression levels of MET associated genes. Fold changes were calculated based on the DESeq2 gene counts in the GATA3-expressed (12 hours after DOX treatment) control or CHD4 knockdown cells compared to DOX minus condition. (d) Bar graphs showing the relative wound closure in the wound healing assay. The wound healing assay was performed in the control or CHD4 knockdown cells before and after GATA3 expression (16 hours). The average values are shown with SDs (N=18, 3 biological replicates x 6 technical replicates).

CHD4 depletion also impacted the biological outcome of GATA3-mediated cell reprogramming, mesenchymal to epithelial transition. MET-related gene expression was exacerbated in the CHD4 knockdown cells (Figure 6c), which is consistent with the increased ATAC-seq signals at newly accessible peaks after CHD4 knockdown. Cell migration assays indicate that GATA3 expression in the CHD4 knockdown cells still showed MET phenotypes at the cellular level, but the degree of cell migration was modestly impacted by CHD4 knockdown (Figure 6d). These results suggest that CHD4 acts to constrain the ability of GATA3 to bind its motif and elicit alterations in gene expression and cellular phenotype.

### CHD4 promotes nucleosome formation over transcription factor motifs

We previously reported that nucleosome remodeling patterns are associated with the chromatin opening activities of GATA3 ^25, 31^. At loci that become accessible following GATA3 induction (newly accessible sites), nucleosome depletion was observed at the center of GATA3 binding peaks, while nucleosome repositioning and accumulation were observed at the constitutively inaccessible GATA3 binding peaks. To understand the impact of CHD4 depletion on nucleosome remodeling following induction of GATA3, we performed MNase-seq to map nucleosomes during the GATA3-mediated cellular reprogramming. Newly accessible peaks exhibit the characteristic pattern of MNase-resistant, positioned nucleosomes flanking a moderately resistant region centered over the transcription factor binding motif (Figure 7a, left panel). When we deplete CHD4, the pattern observed remains unchanged while the amplitude of nuclease resistant nucleosomal peaks flanking the binding site increases. This result suggests that a larger number of alleles within the population sampled have productively bound GATA3, creating a phased array of nucleosomes. To gain further insight into GATA3-induced nucleosome remodeling, we turned to a higher resolution technique, capture MNase-seq, which provides deep mapping of nucleosome positions at individual GATA3 binding loci (Supplementary Figure 5). In this technique, custom designed biotinylated RNA probes are used to enrich defined loci. We designed the RNA probes against ∼2,700 genomic intervals, which contain ∼1800 GATA3 binding sites and ∼940 negative control loci where GATA3 fails to accumulate. Mono-nucleosomal fragment midpoint frequency (dyad frequency) was calculated to monitor nucleosome remodeling during GATA3-mediated reprogramming. In the absence of CHD4, we observed a substantial increase in nucleosomal dyads flanking the GATA motif with no alteration in the final position of the nucleosomes (Figure 7b). These results were consistent with the data from the conventional MNase-seq and suggest that in the absence of CHD4, more alleles within the population sampled exhibit productive binding of GATA3 and create a phased nucleosomal array flanking the binding site. This remodeling is not accompanied by a notable increase in the width of the nucleosome-depleted region.

**Figure 7.**
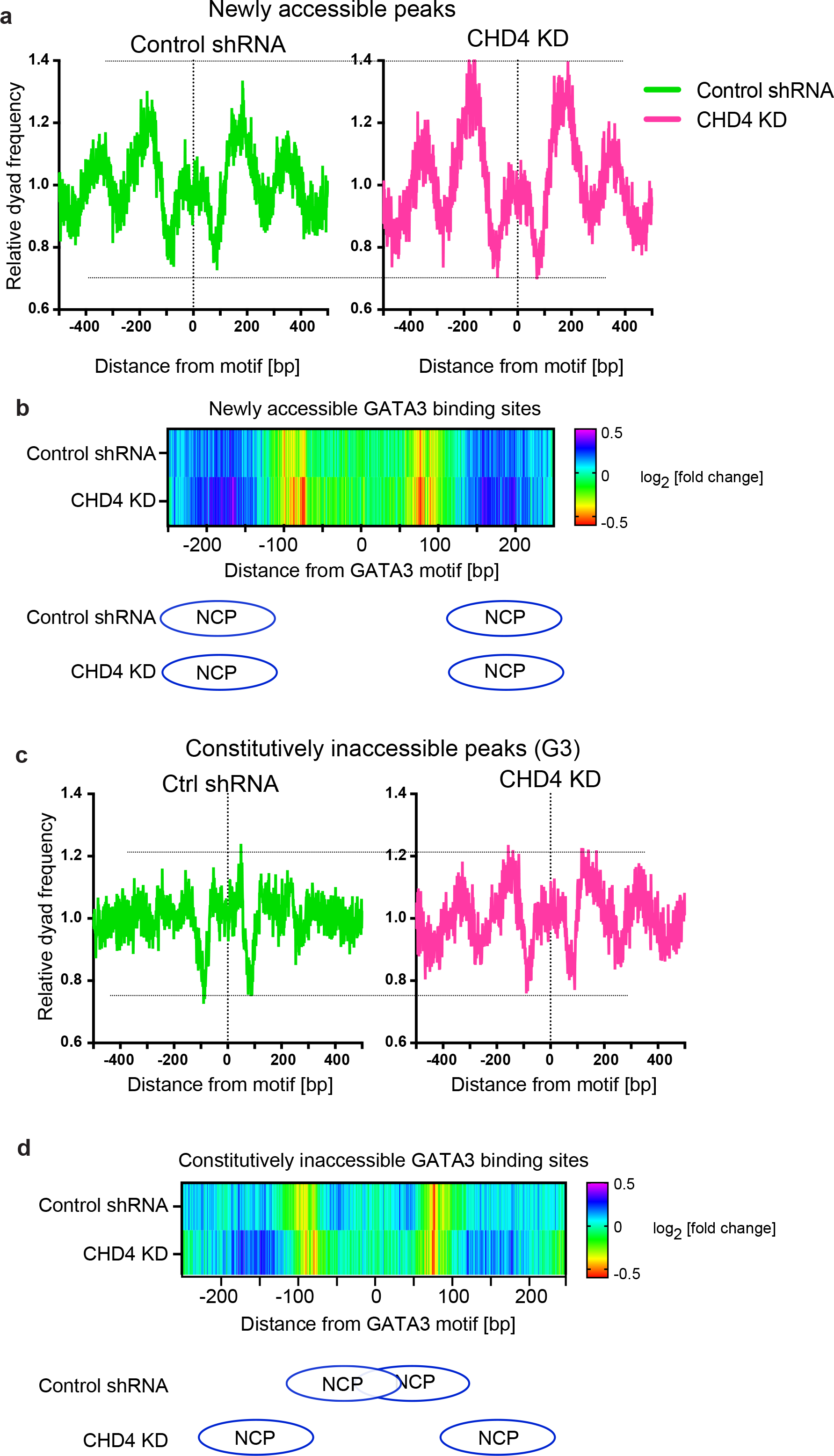
High-resolution nucleosome mapping reveals aberrant nucleosome remodeling induced by CHD4 knockdown **(a)** Metaplot showing averaged dyad frequency at newly accessible peaks in control or CHD4 knockdown cells. The conventional MNase-seq was performed in the control or CHD4 knockdown cells. **(b)** Capture MNase-seq results at the 750 selected newly accessible peaks. Heatmap shows dyad frequency at newly accessible sites relative to time 0 hour. Ellipses indicate the most enriched nucleosome positioning at the GATA3 motif flanking region. **(c)** Metaplot showing averaged dyad frequency at constitutively inaccessible peaks in control or CHD4 knockdown cells. The nucleosome fragments were collected by the conventional MNase-seq. **(d)** Capture MNase-seq data at the 750 selected constitutively inaccessible peaks. Heatmap shows dyad frequency at constitutively inaccessible sites relative to time 0 hour. Ellipses indicate the most enriched nucleosome positions in control (top) or CHD4 knockdown (bottom) cells.

At constitutively inaccessible GATA3 binding sites, we observed a completely different outcome. In conventional MNase-seq, these GATA3-bound loci do not exhibit the phased nucleosomes flanking the binding site that are evident in newly accessible peaks. However, upon CHD4 depletion, we observe a clear pattern of phased nucleosomes flanking the GATA3 binding site that resembles the pattern observed in newly accessible sites (Figure 7c). Capture MNase-seq confirms this observation, showing movement of nucleosomal dyads away from sites where the GATA3 motif is located within the confines of the nucleosome to sites distant from the GATA motif (Figure 7d). At these loci, CHD4 regulates the outcome of GATA3 interaction with the chromatin fiber, its loss changes the outcome from GATA3 bound to the surface of a nucleosome to GATA3-mediated nucleosome eviction from the binding site and establishment of a phased array of flanking nucleosomes.

## Discussion

TFs face multiple challenges in establishing new gene regulatory networks during development, in response to physiologic or environmental signals, and during in vitro cellular reprogramming. These proteins must find appropriate recognition motifs within the genome and they also must contend with physical barriers including chromatin. In eukaryotes with large genomes, including humans, TFs with short recognition motifs must find the correct loci, and only the correct loci, amongst the potentially millions of matches to their binding motif. In theory, degenerate hexameric binding motifs, such as the WGATAR consensus binding motif for GATA3 ^32^, should be present about every 500 base pairs. Based on the human refence genome sequence (hg19), more than 7 million loci contain the GATA3 consensus motif. In most cases, TFs like GATA3 are present on the order of thousands to tens of thousands of molecules per nucleus, meaning there are roughly 100-fold more potential binding sites than TFs.

Chromatin represents a first-line barrier to inappropriate binding of TFs, and it presents a barrier in multiple ways. A subset of TFs cannot read DNA sequence and bind productively at all to DNA wrapped around a histone octamer ^1, 33^. Within the context of a nucleosome, the rotational and translational phasing of DNA necessarily obscures some chemical information where it closely approximates the histone octamer surface, making it unavailable for sequence readers ^5^. Biochemical and structural data indicate that location of binding motifs near the center versus near the periphery of a nucleosome has a strong influence on binding and stability ^31, 34, 35^. Higher order structural features of chromatin, including linker histones, assembly into heterochromatin, or partitioning into low contact frequency nuclear compartments are likely to further restrict the available sequence space to be searched by TFs ^36, 37^. Clearly, other factors must contribute to narrowing the search space and increasing the probability that cellular signals will result in activation (or repression) of the correct subset of genes ^38^.

CHD4 is important for maintenance of chromatin architecture and is known to suppress gene expression during tissue development, cell differentiation, and cell reprogramming ^12, 17, 18, 39, 40^. However, genome-wide mapping finds frequent localization of CHD4 at active gene promoters and enhancers ^21, 24, 41, 42^, clouding understanding of mechanistic roles by which CHD4 regulates gene expression. In this study, we investigated the roles of CHD4 in steady state basal breast cancer cells and during mesenchymal-to-epithelial transition (MET). In the steady state, CHD4 depletion resulted in a primarily increased chromatin accessibility leading to abnormal gene expression unrelated to breast cancer cell identity. While the loci with decreased chromatin accessibility by CHD4 depletion are enriched at promoters, increased chromatin accessibility regions are enriched at intergenic regions. Motif analysis at differential peaks suggested that CHD4 modulates chromatin binding of multiple AP1 family members. In fact, when CHD4 was depleted, at least three AP1 family proteins, JUNB, ATF3, and FRA1, were redistributed, and enrichment at consensus motif-containing regions was reduced.

During GATA3-mediated MET cell reprogramming, CHD4 knockdown again resulted in largely chromatin opening. At both newly accessible and constitutively inaccessible GATA3 bound loci we observed evidence for alterations in local nucleosome positioning, consistent with active roles for chromatin remodelers. Further investigation revealed that depletion of the chromatin remodeling enzyme CHD4, a core subunit of the NuRD complex, dramatically altered this outcome leading to new accessibility at previously inaccessible loci and increasing accessibility to transposition at accessible loci. This abnormal chromatin opening, not observed in the presence of CHD4, was associated with altered gene expression and affected cell fate transition at the cellular level. We speculate that our observations reflect a general property attributable to CHD4, and by extension to NuRD complex. NuRD is found with high frequency at enhancers ^21, 24, 41, 42^ where it is integral to the process of enhancer decommissioning during development ^10, 17, 41^ and reprogramming ^39, 40^. We propose that NuRD acts, in part, to antagonize transcription factor driven increases in chromatin accessibility – regardless of the ultimate outcome. It seems plausible that local translational motion of nucleosomes relative to transcription factor binding motifs leads to architectural obstacles to motif recognition. At loci that become accessible, co-binding of multiple TFs or recruitment of other chromatin modification/remodeling enzymes generates a competition between factors promoting and opposing DNA accessibility that ultimately reaches a dynamic equilibrium in which some alleles within the population are accessible to structural probes. At loci that fail to become accessible, failure to recruit activating co-factors leads to generation of a very different type of equilibrium, one in which nucleosome translational position relative to the GATA motif permits GATA3 binding to nucleosomal DNA in the absence of detectable accessibility. In this manner, choice of binding sites within the genome can be ‘proof-read’ by chromatin remodelers simply through enforcing a highly dynamic state ^43, 44^. In principle, such chromatin dynamics would promote two critical outcomes: they would provide assurance against inappropriate transcriptional activation following transcription factor binding at incorrect sites and they would provide an opportunity for rapid decommissioning of enhancers to enable progression to new patterns of gene expression consistent with cellular needs.

## Methods

### Cell line and cell culture

MDA-MB-231 cells were originally obtained from ATCC. MDA-MB-231 cells were cultured in DMEM high-glucose medium with 10% FBS (Thermo Fisher Scientific or R&D Systems). Doxycycline-inducible GATA3 expression system in MDA-MB-231 cells was developed by lentiviral transduction. Ty1-tagged GATA3 gene was inserted into the pINDUCER20 vector, and the lentivirus was generated by the 2nd generation lentiviral plasmids system using 293T cells. pINDUCER20 was a gift from Guang Hu (NIEHS/NIH). psPAX2 and pMD2.G were gifts from Didier Trono (Addgene plasmid #12260, #12259). After antibiotic selection with G418, cell colonies were collected, and GATA3 expression levels were investigated by western blot. The cell clone that has low basal GATA3 expression (without DOX treatment) was used in this study.

### ChIP-seq

ChIP-seq libraries were prepared as previously described ^47^. Briefly, DOX treated cells were fixed at 12 hours with 1% formaldehyde. Fixed cells were treated with hypotonic buffer containing 10 mM HEPES-NaOH pH 7.9, 10 mM KCl, 1.5 mM MgCl2, 340 mM sucrose, 10% glycerol, 0.5% Triton X-100, and protease inhibitor cocktail (Thermo Fisher Scientific). Chromatin was digested by sonication with Covaris S220 in the lysis buffer containing 20 mM Tris-HCl pH 8.0, 2 mM EDTA, 0.5 mM EGTA, 0.5 mM PMSF, 5mM sodium butyrate, 0.1% SDS, and protease inhibitor cocktail. 2.5 µg of each antibody was added to each chromatin solution (1 million cells/reaction). After overnight incubation, protein A/G mixed Dynabeads were added, and the samples were rotated for 2 hours. Eluted DNAs were reverse crosslinked at 65 °C for 4 hours, followed by the incubation with proteinase K for 1 hour and purified by AMPure XP (Beckman Coulter).

ChIP-seq libraries were generated by the NEXTflex Rapid DNA-seq kit (PerkinElmer) and sequenced on NextSeq 500 (Illumina, paired-end) at the NIEHS Epigenomics Core Facility. The same data processing protocol to ATAC-seq was used. Mapped reads were converted to a single fragment and used to generate genome coverage tracks on the UCSC Genome Browser and for metaplot analyses.

To carry out correlation and differential analysis, we used read counts from the reference peak sets and processed the data using the SARTools pipeline ^48^ with edgeR.

### ATAC-seq

The pGIPZ vectors and lentiviruses encoding CHD4 shRNA and control shRNA were provided by the NIEHS Epigenomics Core and Viral Vector Core. 400,000 cells were plated on 6 cm dishes, and infected with these shRNA lentiviruses at 24 hours and 32 hours after plating. After overnight incubation, the medium was replaced with 4 ml of fresh medium and further incubated for 2 days. The GATA3 expression was initiated by adding DOX (1 µg/ml at final) in fresh medium. 12 hours after induction, cells were harvested and resuspended in PBS(-).

The ATAC-seq libraries were prepared as previously described ^26, 47^. 25,000 cells (in 25 ul) were transferred to new tubes, and nuclei were isolated with CSK buffer (10 mM PIPES pH 6.8, 100 mM NaCl, 300 mM sucrose, 3 mM MgCl2, 0.1% Triton X-100).

Nuclei were treated with 2.5 µl of Tn5 Transposase (Illumina) in the standard tagmentation reaction buffer (25 µl). The total of 8 PCR cycles were performed to amplify the DNA fragments, and the libraries were sequenced on NextSeq 500 at the NIEHS Epigenomics Core Facility. The raw sequence reads were filtered based on the mean base quality score >20. Adapter sequences were removed by Trim Galorre! (Babraham Institute). Processed reads were mapped to hg19 genome using Bowtie 0.12.8 ^49^, and uniquely mapped reads (non-duplicate reads) were used for the subsequent analysis.

Peak classification was conducted based on the following criteria.

Newly accessible peaks are defined as the GATA3 peaks that show (1) >2-fold increase in ATAC-seq signals at GATA3 peaks (+/- 200 bp from peak center) compared to the control (Time 0h) condition and (2) >30 normalized reads at the peak flanking (+/- 1 kbp) region.

Constitutively accessible peaks are defined as the GATA3 peaks that show >30 normalized reads at the flanking regions (+/- 1 kbp).

Constitutively inaccessible peaks are defined as the GATA3 peaks that show <30 normalized reads at the flanking regions.

### Capture MNase-seq

Capture MNase-seq was performed as previously described ^31^. Biotinylated RNA probes were designed and purchased via SureSelect Custom DNA Target Enrichment Probes system with the following probe design specification (Tiling density 2x, Masking: Least Stringent, Boosting: Balanced). The target regions are listed on Supplementary table 1.

Nucleosomal fragments were prepared by digesting nuclei with Micrococcal nuclease (MNase). Sequencing libraries were prepared by NEXTflex Rapid DNA-Seq kit (PerkinElmer). Libraries from different conditions were pooled, and 750 ng DNAs were used to perform nucleosomal DNA fragment enrichment at a subset of GATA3 peaks with the SureSelect Enrichment kit. After the fragment enrichment by RNA probe hybridization and streptavidin pull-down, captured DNAs were amplified by PCR (12 cycles) and sequenced on NextSeq 500 at the NIEHS Epigenomics Core Facility.

The same data processing protocol to ATA-seq was used for the capture MNase-seq data but duplicate reads were retained. To generate heatmaps, midpoints only from mono-nucleosomal fragments (120-170 bp) were collected. The midpoint frequency (considered as dyad frequency) was normalized by the data from Time 0.

### RNA-seq

Total RNAs were purified by the Qiagen RNeasy kit. Sequencing libraries were generated by TruSeq RNA library preparation kit with Ribo-Zero and sequenced on NextSeq 500 (Illumina) at the NIEHS Epigenomics Core Facility. Filtered read pairs were mapped to hg19 by STAR (version 2.5) with the following parameters: -- outSAMattrIHstart 0 --outFilterType BySJout --alignSJoverhangMin 8 -- out- MultimapperOrder Random ^50^. Subread featureCounts (version 1.5.0-0-p1) and DESeq2 (v1.10.1) with FDR < 0.01 were used to define differentially expressed genes (DEGs) ^51^. Pathway analysis was performed by DAVID database ^45, 46^ using DEGs that have |log2 (fold change)| >0.5. DEGs are summarized in Supplementary Table 2.

### Migration assay

The control shRNA or CHD4 shRNA expressing MDA-MB-231 cells were prepared as described above using the lentiviruses. The GATA3-induced cells were culture for at least 2 days. 4 million were seeded in each 6-well plate and grown overnight. The confluent cells were scratched with a 1 ml micropipette tip. After scratching, the cells were washed with 2 ml of pre-warmed serum-free medium, and the same volume of the serum-free medium was added to each well. Cell images were taken at multiple time points (0, 8, 16, 24, and 33 hours) by Olympus IX71 microscope, and imageJ (Version: 2.1.0/1.53c) was used to quantify wound areas.

### Antibodies

Anti-Ty1 antibody (MilliporeSigma, Imprint monoclonal BB2), JUNB (Cell Signaling, C37F9), ATF3 (Cell Signaling, E9J4N), FRA1 (Cell Signaling, D80B4), and CHD4 Abcam (Abcam, ab72418) was used for ChIP-seq and western blot. GATA3 antibody was generated in rabbits using recombinant 6x histidine tag-fused GATA3 full-length wild-type protein ^52^.

### Data availability

Data generated in this study are available at Gene Expression Omnibus under Accession Number GSE201797, GSE201798, GSE201799, GSE201800, and GSE201801. GATA3 ChIP-seq data in the stable cell line (GSE72141) were previously generated ^25^.

## Supporting information

CaptureMNase-seq_Probes

RNA-seq_DEGs

## Acknowledgments

We gratefully acknowledge the UND Genomics core facility and the NIEHS/NIH core facilities (Epigenomics Core, Viral Vector Core, Integrated Bioinformatics Core) for outstanding technical assistance. We thank Dr. Archana Dhasarathy for help with cell migration assay, Dr. Negin Martin for help with lentivirus preparation, and Mr. Michael Hill and Dr. Bony De Kumar for help with next generation sequencing analysis. We thank Drs. Sergei Nechaev and Archana Dhasarathy for insightful suggestions on the manuscript.

---

**Supplementary Figure 1.**
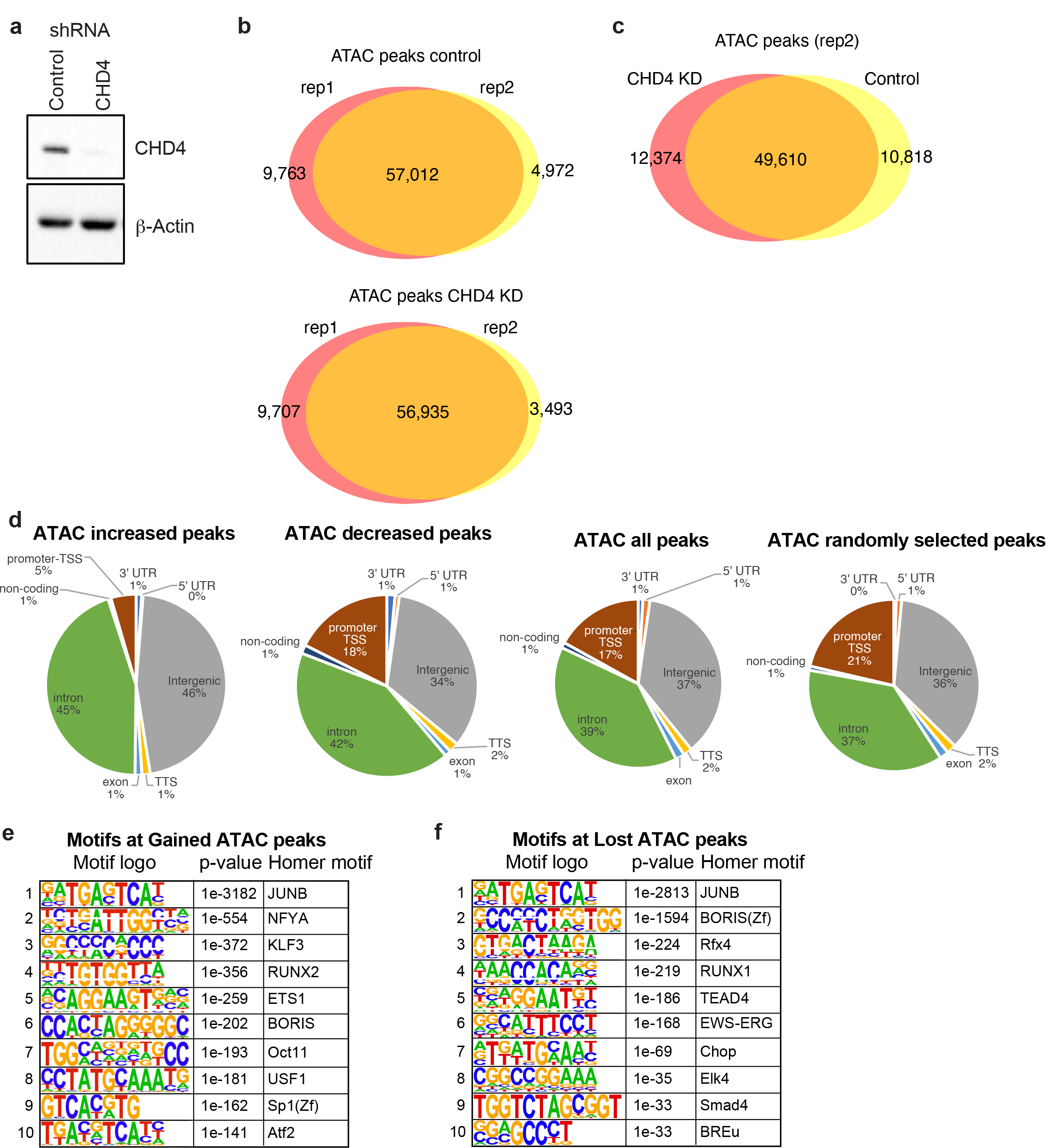
ATAC-seq peak overlap in CHD4 knockdown cells (a) Western blot showing CHD4 knockdown. MDA-MB-231 cells were infected with the lentivirus encoding control shRNA or CHD4 shRNA. β-Actin expression was used as an internal control. **(b)** Venn diagram showing the ATAC-seq peak overlap between biological replicates in control (top) or CHD4 knockdown (KD, bottom) cells. **(c)** Venn diagram showing the ATAC-seq peak overlap between control and CHD4 knockdown MDA-MB-231 cells. **(d)** Pie chart showing peak annotation defined by HOMER. Increased, decreased, all, or randomly selected ATAC-seq peaks are classified into 8 annotation categories. **(e-f)** HOMER de novo motif analysis. ATAC-seq peaks are grouped in Gained (e, uniquely observed in CHD4 knockdown cells) or Lost (f, only observed in the control cells) ATAC-seq peaks (shown in Figure 1b), and are used as input for HOMER motif analysis.

**Supplementary Figure 2.**
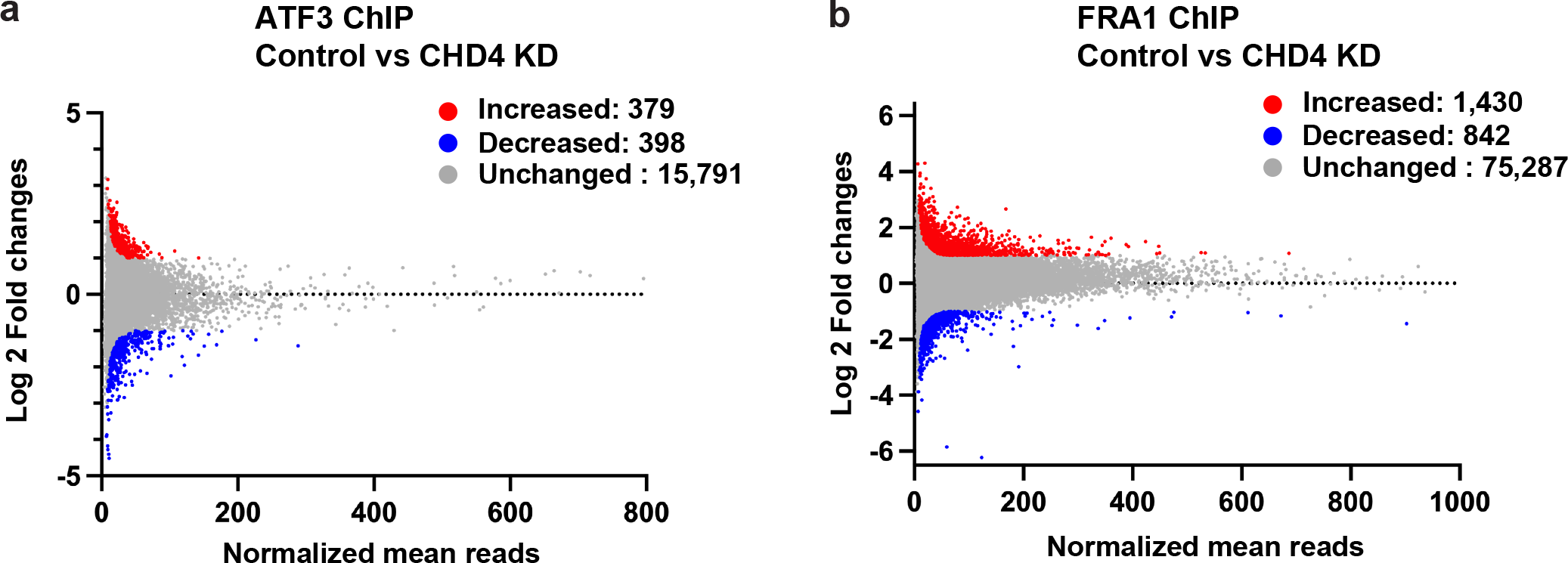
Differential peak analyses of ATF3 and FRA1 ChIP-seq data. (a) Scatter plot shows increased (red), decreased (blue), and unchanged (grey) ATF3 peaks upon CHD4 depletion. **(b)** Scatter plot shows increased (red), decreased (blue), and unchanged (grey) FRA1 peaks upon CHD4 depletion.

**Supplementary Figure 3.**
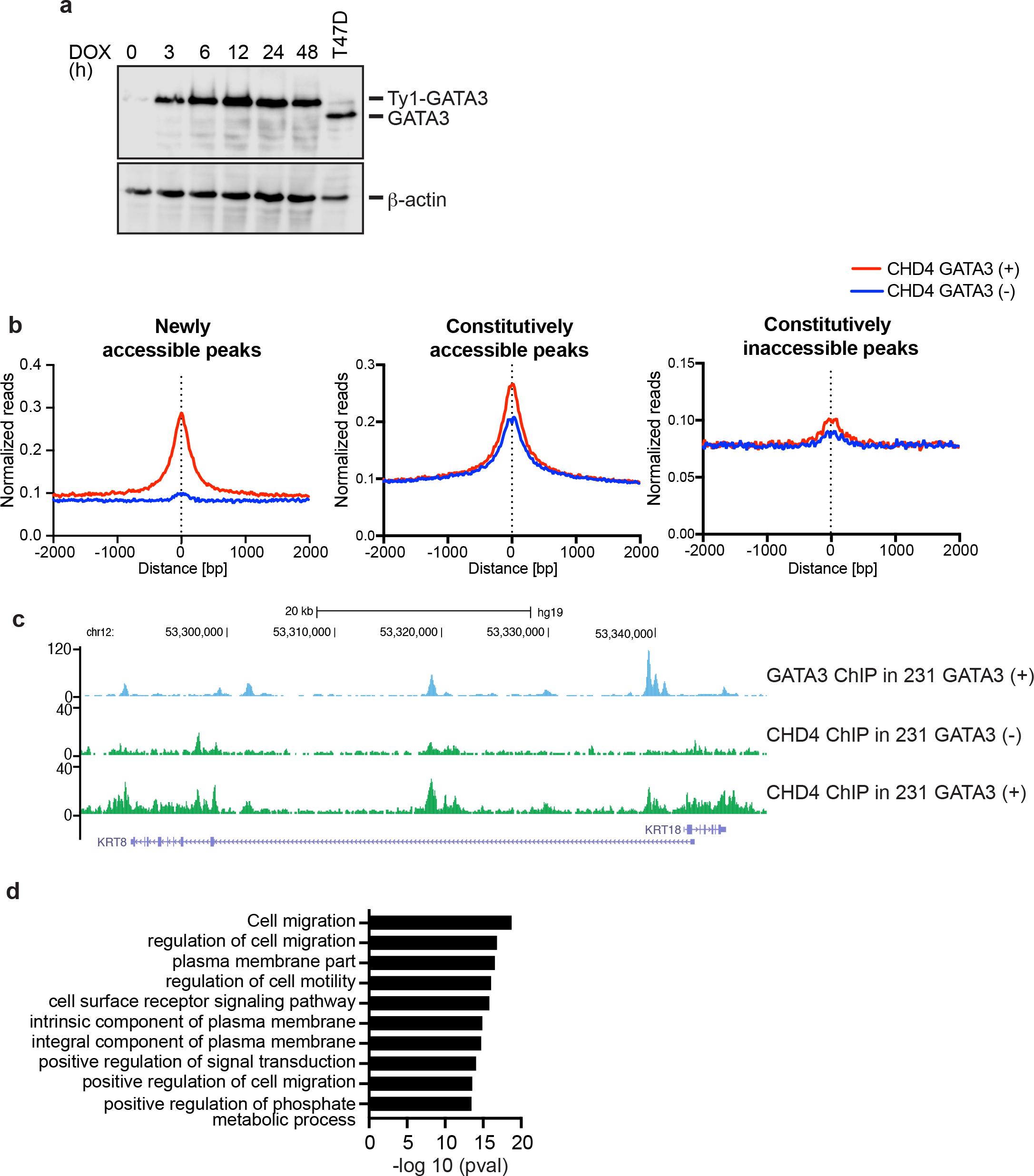
De novo motif analysis of GATA3 peaks (a) Western blot showing GATA3 expression after DOX treatment (1 μg/ml at the final concentration). β-Actin expression was used as an internal control. **(b)** Metaplots showing the CHD4 ChIP-seq signals before and after GATA3 expression. Newly accessible, constitutively accessible. constitutively inaccessible peak groups are previously defined in the GATA3 stably expressed cell system 25. **(c)** Representative genome track of GATA3 and CHD4 ChIP-seq data. GATA3 ChIP-seq was performed in the GATA3-expressed stable cell line. CHD4 ChIP-seq was performed in the GATA3 negative or positive MDA-MB-231 stable cell lines. **(d)** Pathway analysis of the up-regulated genes shown in Figure 4a. Top 10 enriched pathways are shown. expression system.

**Supplementary Figure 4.**
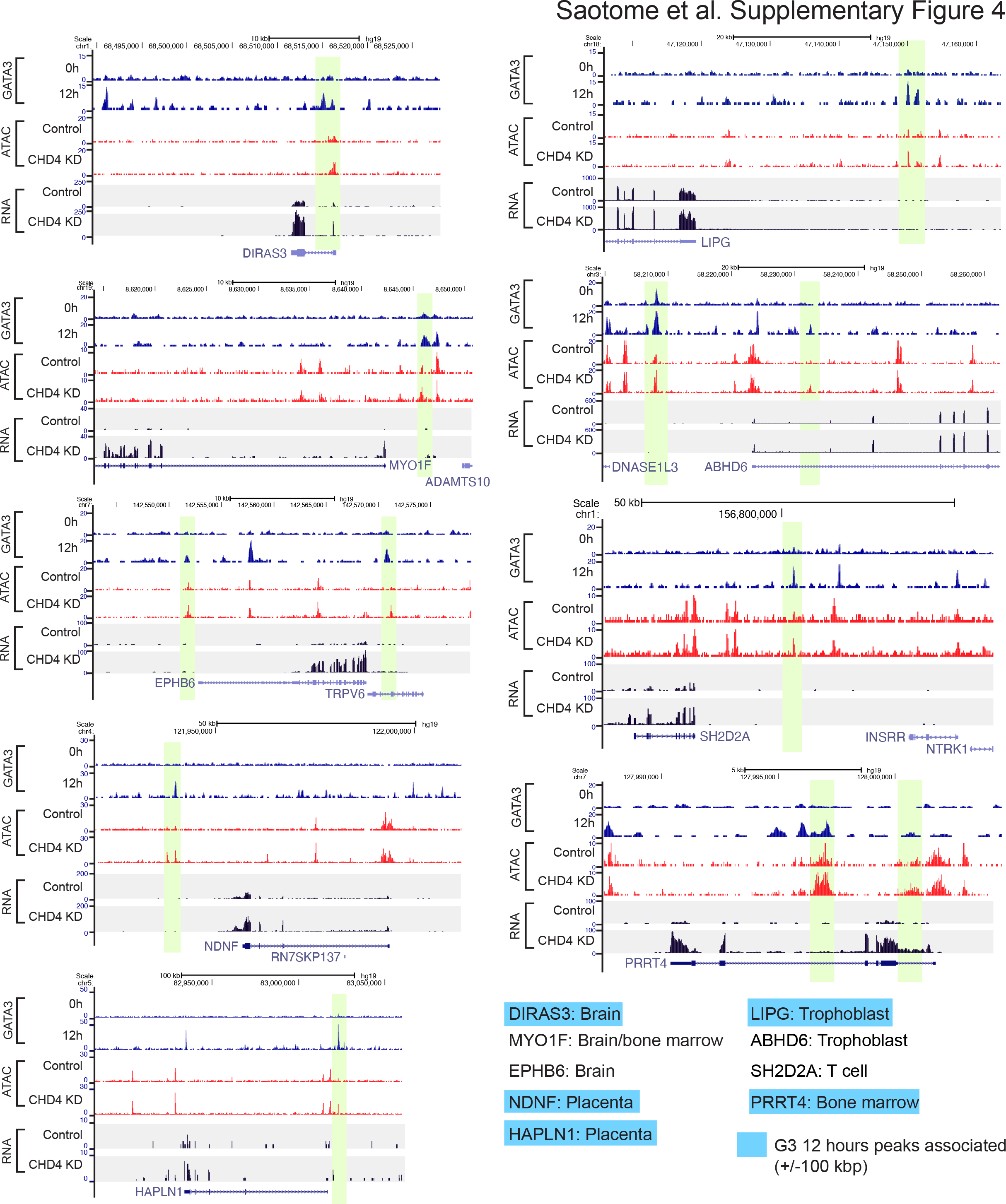
CHD4 depletion induces aberrant chromatin opening and gene activation. Genome browser tracks show the examples of aberrant gene expression. Brain, placenta, trophoblast, T cell, and bone marrow related genes are selected. In each figure, GATA3 peaks that have increased ATAC-seq signals in the CHD4 knockdown cells are highlighted. The genes that are associated with the constitutively inaccessible GATA3 peaks (peaks within +/- 100 kbp) were highlighted in light blue.

**Supplementary Figure 5.**
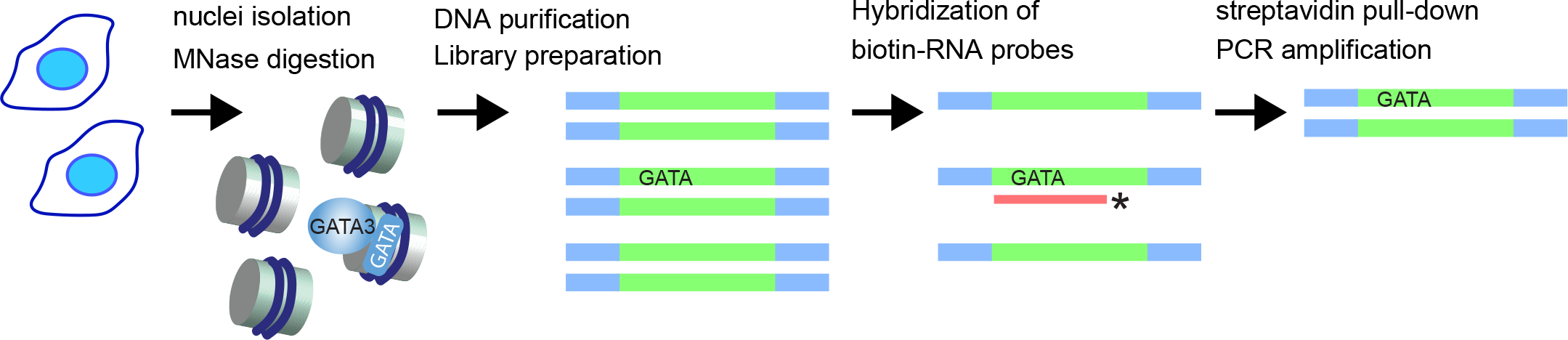
Capture MNase-seq. Experimental scheme of capture MNase-seq. Mono-nucleosmal fragments were prepared by MNase digestion. Sequencing libraries were made by NEXTFLEX Rapid DNA-Seq kit (PerkinElmer). Biotinylated RNA probes (Agilent) were used to enrich nucleosome fragments at selected GATA3 peaks.

